# Characterization of amyloid β fibril formation under microgravity conditions

**DOI:** 10.1101/2020.03.23.004341

**Authors:** Maho Yagi-Utsumi, Saeko Yanaka, Chihong Song, Tadashi Satoh, Chiaki Yamazaki, Haruo Kasahara, Toru Shimazu, Kazuyoshi Murata, Koichi Kato

## Abstract

Amyloid fibrils are self-assembled and ordered proteinaceous supramolecules structurally characterized by the cross-β spine. Amyloid formation is known to be related to various diseases typified by neurogenerative disorders and involved in a variety of functional roles. Whereas common mechanisms for amyloid formation have been postulated across diverse systems, the mesoscopic morphology of the fibrils is significantly affected by the type of solution condition in which it grows. Amyloid formation is also thought to share a phenomenological similarity with protein crystallization. While many studies have demonstrated the effect of gravity on protein crystallization, its effect on amyloid formation has not been reported. In this study, we conducted an experiment at the International Space Station (ISS) to characterize fibril formation of 40-residue amyloid β (Aβ(1-40)) under microgravity conditions. Our comparative analyses revealed that the Aβ(1-40) fibrilization progresses much more slowly on the ISS than on the ground, similarly to protein crystallization. Furthermore, microgravity promoted the formation of distinct morphologies of Aβ(1-40) fibrils. Our findings demonstrate that the ISS provides an ideal experimental environment for detailed investigations of amyloid formation mechanisms by eliminating the conventionally uncontrollable factors derived from gravity.

## Introduction

Certain proteins are known to self-assemble into ordered supramolecular structures, such as filaments and even crystals, under specific physiological and pathological conditions^1,2^. The cytoskeletons present in all cells are made of such filamentous proteins, which dynamically assemble and disassemble to control cell morphology, movement, and signaling in physiological processes^1,3^. In contrast, filamentous protein aggregates known as amyloid fibrils are actively involved in pathological processes and are associated with various diseases, including neurogenerative disorders and diabetes^4–6^. Each of these diseases is characterized by a specific amyloidogenic protein, but it has been suggested that a common molecular mechanism governs fibril formation across these diverse systems. Amyloids may also play a variety of functional roles in many organisms and have been regarded as potentially useful nanomaterials in recent years^7–9^. Therefore, a detailed characterization of the self-assembling mechanism of amyloid fibrils and the resulting morphology is essential for developing therapeutic strategies for amyloidopathy and gaining the knowledge needed to create future nanomaterials.

Increasing evidence provided by cryo-electron microscopy (cryo-EM)^10–12^ and solid-state NMR spectroscopy^13–15^ demonstrate that the morphology of amyloid fibrils is significantly affected by various solution conditions, such as protein concentration, ionic strength, pH, temperature, and pressure^9^. X-ray diffraction studies show that amyloid fibrils share similar structural features characterized by a cross-β spine: a double β-sheet with each sheet running parallel to the fibril axis. At the mesoscopic level, however, amyloid fibrils formed under the same conditions show considerable morphological diversity^10,11^. These molecular polymorphisms are supposed to be derived from differences in the number, relative orientation, and internal substructure of the protofilaments. Despite the cumulative structural data, a comprehensive understanding of the molecular mechanisms behind amyloid polymorphisms remains largely unexplored, as a variety of factors can influence the molecular assembly process.

Recent evidence suggests that amyloid formation and protein crystallization share phenomenological similarities during molecular assembly processes: both protein crystals and amyloid fibrils can form in a supersaturated solution via initial nucleation and subsequent growth despite their morphological differences^16^. In the case of protein crystallization, gravity notably influences the assembly process, as it causes sedimentation and convection flow, thereby perturbing microenvironments surrounding the crystal^17,18^. Hence, protein crystals grown under microgravity conditions often exhibit increased size, integrity, and internal order. Such microgravity-grown crystals have been appreciated in crystallographic analyses because of their improved quality of diffraction to a higher resolution. In contrast, there is no report on the effect of gravity on amyloid fibril formation. In this study, we conducted the “Amyloid” experiment at the International Space Station (ISS) to characterize amyloid formation under a microgravity environment. We focused on amyloid β (Aβ) proteins, which are 40- or 42-amino-acid peptides cleaved from its precursor membrane protein by β- and γ-secretases^19^. The conversion of soluble monomeric Aβ to aggregated toxic form, such as oligomers and fibrils, is a crucial step in the development of Alzheimer’s disease^19,20^. EM studies have shown that Aβ amyloid fibrils exhibit significant structural heterogeneity^10,11,21–26^ (**Figure 1**). We compared the kinetics of Aβ fibrilization and the resulting molecular morphology under microgravity and on-the-ground conditions.

**Figure 1:**
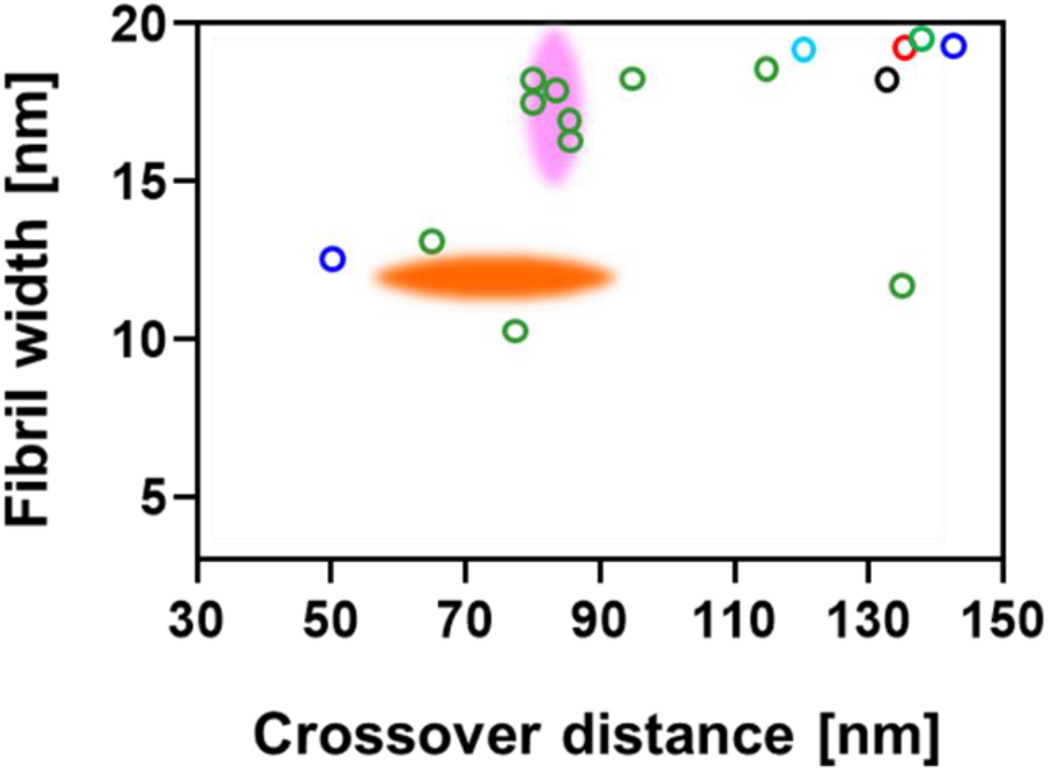
Polymorphism of the Aβ(1-40) amyloid fibrils. Distributions of crossover distance and the widest fibril width for the Aβ(1-40) amyloid fibrils previously reported. Aβ(1-40) amyloid fibrils prepared in 50 mM sodium borate, pH 7.8 at 4°C^22^ (black), in 50 mM sodium borate, pH 7.8 at 22°C^23^ (green and pink), in 50 mM sodium borate, pH 8.7 at 4°C ^25^ (cyan), in 50 mM sodium borate, pH 9.0 at 20°C ^21^ (red), in 50 mM Tris-HCl, pH 7.4 at room temperature^24^ (blue), in PBS, pH 7.4 at 37°C ^23^ (orange).

## Results

### Amyloid formation under microgravity

A microgravity experiment on amyloid formation was conducted at the Japanese Experiment Module, KIBO, on the ISS from December 16, 2017 to January 14, 2018. Four sets of frozen samples of Aβ(1-40) solution were transferred to the ISS and stored at −95°C. All samples were simultaneously thawed by incubation at 2°C for 16 h. Subsequently, the samples were immediately transferred to an incubator set at 37°C and continuously incubated at 37°C to promote amyloid fibril formation. The recorded temperatures were downlinked at constant intervals to the Tsukuba Space Center (Tsukuba, Japan). The four sets of samples were transferred to −95°C cold stowage at different time intervals: 6 hours, 1, 3, and 9 days after incubation at 37°C (**Figure 2**). All sets of the samples were stored at −95°C until they were returned to ExCELLS (Okazaki, Japan) after 10 days and kept at −80°C until used for experimental observations. All experimental procedures were performed smoothly on ISS-KIBO without accidents. Control sample sets were independently processed under the same conditions on the ground.

**Figure 2:**
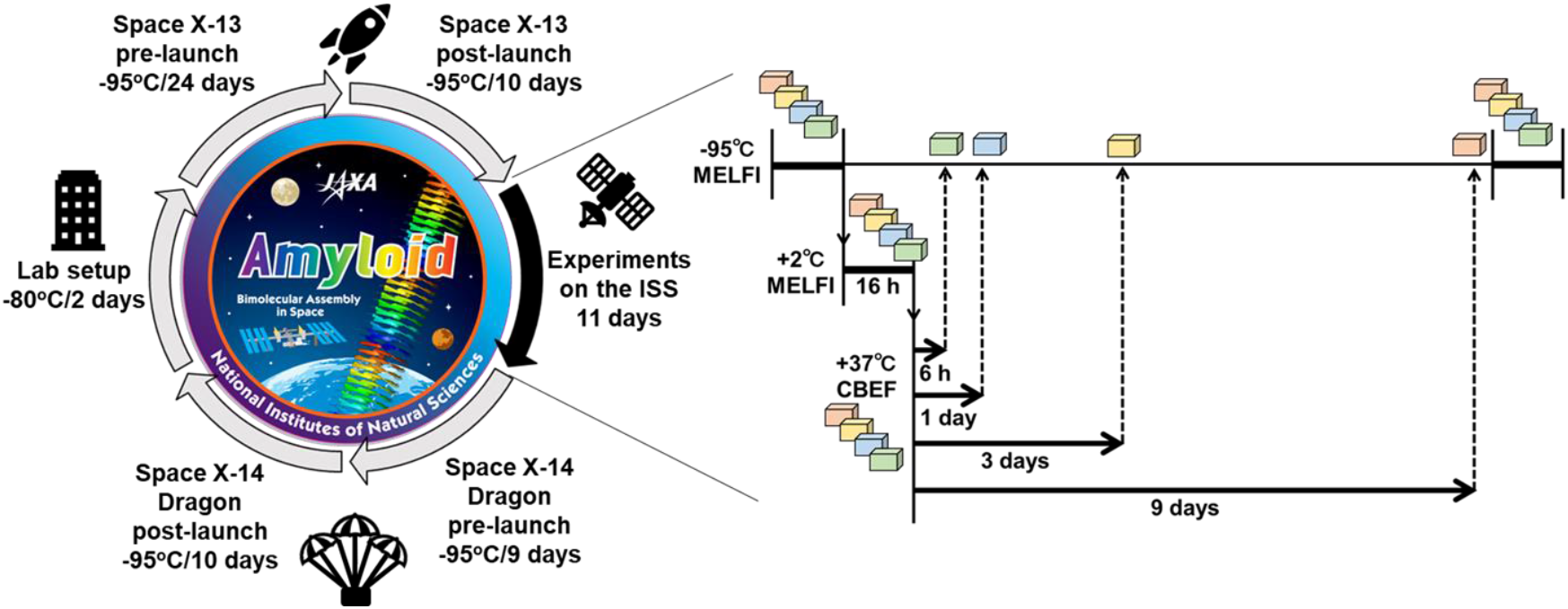
Timeline and procedure of the microgravity experiments performed on the ISS.

### Effect of microgravity on the amyloid formation

To investigate the effect of the microgravity environment on the macroscopic steps underlying the amyloid formation process of Aβ(1-40), a global analysis of the aggregation profiles of Aβ(1-40) incubated on the ISS and on the ground was performed using the amyloid-binding dye thioflavin T (ThT) as a fluorescent probe. As shown in **Figure 3**, Aβ(1-40) incubated on the ISS formed ThT-reactive species at a much slower rate than that incubated on the ground. While the increase in fluorescence reached a plateau after 3 days on the ground, it did not reach a plateau after 9 days under microgravity. Despite the limited data points, the fluorescence growth apparently exhibited a longer lag time before the rise and a more gradual increase, suggesting that microgravity slowed down both the nucleation and elongation steps.

**Figure 3:**
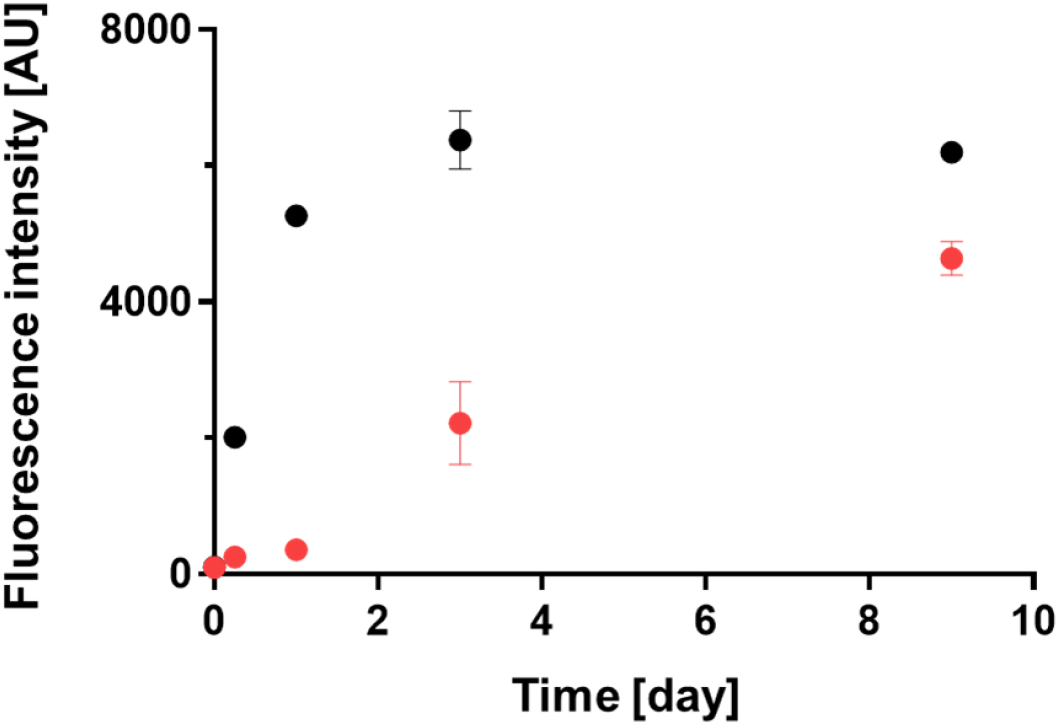
Amyloid formation process of Aβ(1-40). Aβ(1-40) solutions were incubated at a concentration of 100 μM under microgravity (red) and on the ground (black). Each intensity value is the mean ± SD of the three values.

### Effect of microgravity on amyloid morphology

To investigate the potential effects of the microgravity environment on amyloid fibril structure, Aβ(1-40) fibril morphology between the microgravity-grown fibrils and the ground-grown fibrils was compared using cryo-EM combined with 3D image reconstruction. The cryo-EM data revealed that the Aβ(1-40) solution incubated for 9 days under microgravity conditions contained well-separated, unbranched fibrils with periodic structures. Despite considerable morphological variations, the fibrils were found to be classified into two distinct types (termed hereinafter Type-1 and Type-2) in terms of fibril width at the crossovers (white arrows in **Figure 4**). Type-1 fibrils, which made up about 80% of total fibrils, exhibited narrow widths (6-8 nm) at the crossover points, while Type-2 fibrils, which accounted for the remaining 20%, had broader widths (>10 nm) at the crossover points. Each type of fibril was further subdivided based on the widest fibril width (yellow arrows in **Figure 4**) between two crossovers and the crossover distance. Statistical analysis of the fibril morphology between Type-1 and Type-2 fibrils indicated that these two fibril types were essentially indistinguishable in fibril width and crossover distance, despite a significant distribution shift regarding these geometrical parameters: The ranges of fibril width and crossover distance were 3.8-22.5 nm and 39.3-148.7 nm for Type-1 and 6.8-17.2 nm and 43.9-144.5 nm for Type-2, respectively (**Figure 5**). The averaged fibril width and crossover distance were 9.2±0.1 nm and 73.4±0.9 nm for Type-1 and 11.7±0.2 nm and 74.7±1.7 nm for Type-2, respectively.

**Figure 4:**
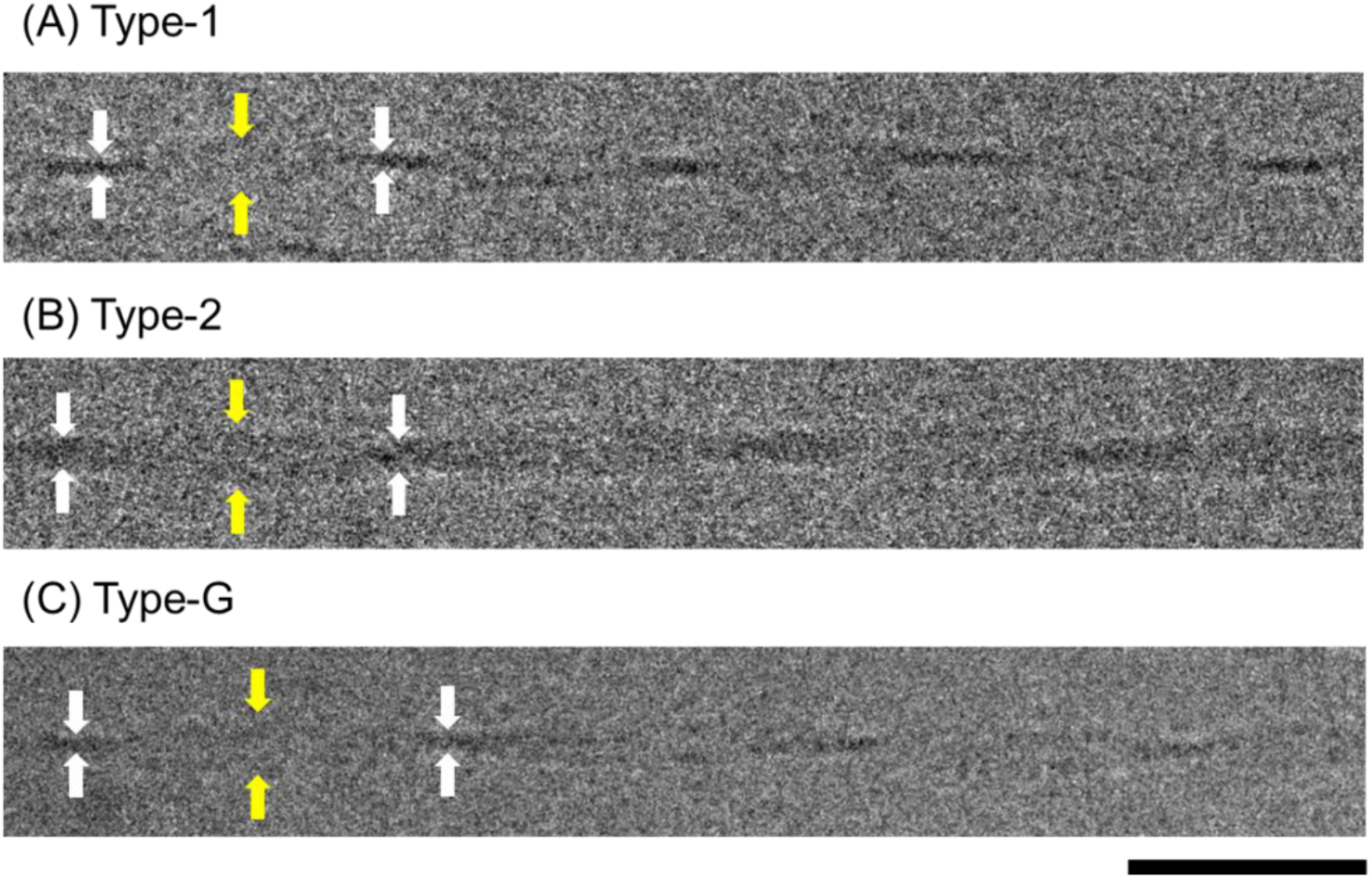
Cryo-EM images of the Aβ(1-40) amyloid fibrils. Representative electron micrographs of (A) Type-1, (B) Type-2, and (C) Type-G fibrils. Crossovers and the widest points between crossovers are indicated by white and yellow arrows, respectively. Scale bar is 50 nm.

**Figure 5:**
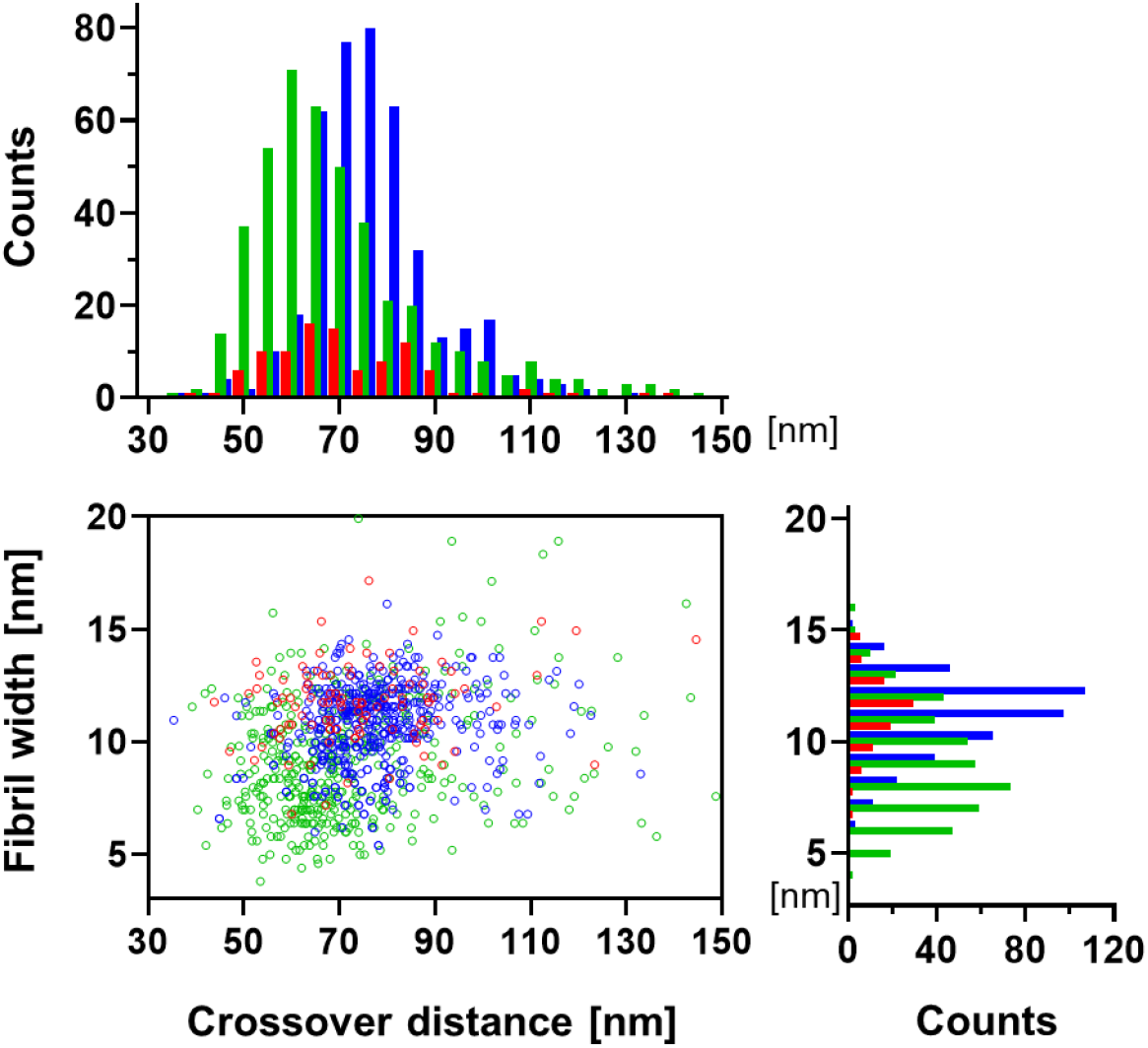
Distribution of crossover distances and fibril widths (measured at the widest points between crossovers) for all three types of fibrils. Total counts of Type-1 (green), Type-2 (red), and Type-G (blue) were 433, 98, and 410, respectively. The images were analyzed using the Fiji software^36^.

In contrast, the Aβ(1-40) fibrils prepared on the ground belonged only to morphology similar to Type-1 with variations in the fibril width (10.7±0.1 nm) and crossover distance (77.7±0.6 nm) : The ranges of fibril width and crossover distance of these ground-grown Aβ(1-40) fibrils were 5.4-16.1 nm and 35.4-133.0 nm, respectively (**Figures 4 and 5**). It should be noted that the distribution of these parameters was significantly shifted to larger values in the ground-grown Aβ(1-40) fibrils compared to the microgravity-grown Type-1 fibrils. Hereinafter, the morphological type of the Aβ(1-40) fibrils grown on the ground is referred to as Type-G.

### 3D reconstruction of amyloid fibrils

The 3D densities of typical amyloid fibrils with the highest incidence in Type-1, Type-2, and Type-G were reconstructed on the basis of the cryo-EM data (**Figure 6, Table 1**). In each case, the 3D-reconstruction was performed helically, assuming 2-fold rotational symmetry around the fibril axis based on the raw image. Table 1 summarizes the details of the reconstruction and the properties of the reconstructed fibrils. The averaged 3D structures of Type-1 and Type-2 fibrils were 10.3 nm and 11.6 nm width, the crossover pitches were 68.0 nm and 65.0 nm, and the cross-section areas were 126.7 nm^2^ and 277.3 nm^2^, respectively. Type-G fibrils exhibited a width of 11.2 nm, a crossover pitch of 75.0 nm, and a cross-section area of 153.0 nm^2^. Atomic force microscopy showed that all helical fibrils possessed a left-handed chirality. The cross-sectional structures of Type-1 and Type-G fibrils exhibited a compact shape, while the cross-section of Type-2 accommodated a square trace with symmetrical orientation, suggesting their distinct differences in the relative orientation and internal substructure of protofilaments.

**Figure 6:**
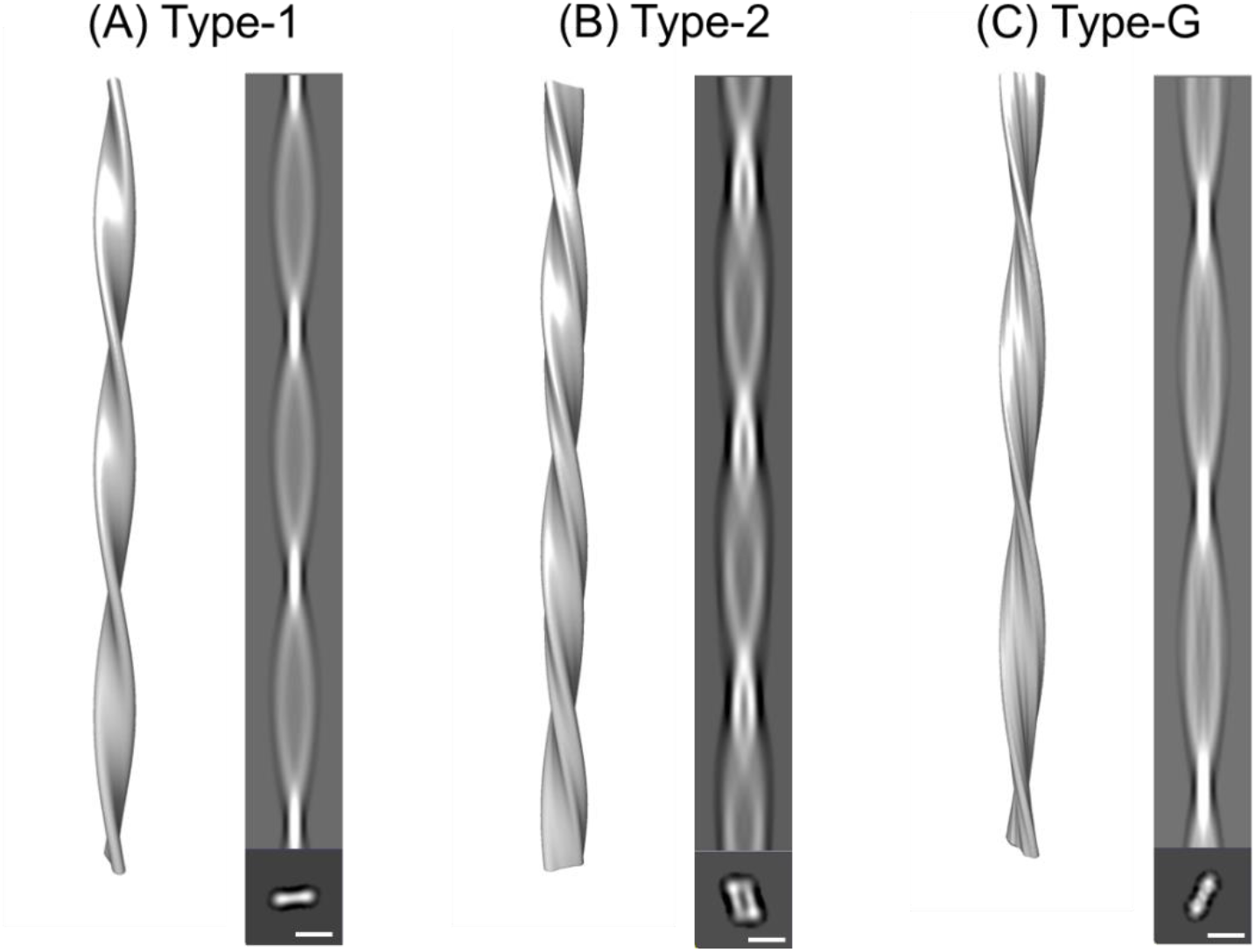
3D image reconstruction of Aβ(1-40) amyloid fibrils. Side-view surface rendering of the fibril reconstruction (left), the reprojection (right, upper) and cross-section (right, lower) of the 3D reconstruction of (A) Type-1, (B) Type-2, and (C) Type-G. Scale bar is 10 nm.

**Table 1:** Reconstruction details and properties of the reconstructed fibrils

|  | Type-1 | Type-2 | Type-G |
| --- | --- | --- | --- |
| Micrographs | 29 | 12 | 23 |
| Segment number | 1452 | 404 | 774 |
| Scale on specimens ($\text{\AA}/\text{pix}$) | 3.984 | 3.984 | 3.984 |
| Box size (nm) | 220 | 220 | 220 |
| Helical rise ($\text{\AA}$) | 4.76 | 4.76 | 4.76 |
| Crossover distance (nm) | 68.0 | 65.0 | 75.0 |
| Fibril width (nm) | 10.3 | 11.6 | 11.2 |
| Cross-sectional area ($\text{nm}^2$) | 126.69 | 277.28 | 152.98 |
| Resolution ($\text{\AA}$) | 32 | 33 | 17 |

## Discussion

In this study, we found significant differences in amyloid formation kinetics and fibril morphology between microgravity-grown and ground-grown Aβ(1-40) amyloids. To the best of our knowledge, this is the world’s first observation of the effect of gravity on amyloid formation. Under microgravity conditions, Aβ(1-40) fibrilization occurred much more slowly than on Earth (**Figure 3**). Intriguingly, a similar trend has been observed in protein crystallization, which often occurs more slowly under microgravity conditions than on the ground. Such a delay in crystallization has been primarily associated with the lack of convection effects under microgravity conditions^18^. In this circumstance, the diffusion rate of macromolecules is so slow that a concentration gradient is generated, giving rise to protein depletion zones around the nucleus and growing crystals even under supersaturation conditions. This causes a convection flow in the presence of gravity, agitating rapid crystal growth, though accompanied with lower orders and more defects. However, microgravity eliminates such convective mixing, often resulting in very slow crystal growth^17,27–29^. In addition, the protein depletion zone formed under microgravity can serve as a filter against larger impurities with much slower diffusion, such as protein aggregates, which potentially bind to the crystal surface and impair the quality of crystal^30,31^. Therefore, the lack of convection mixing is advantageous in improving crystal integrity and size in microgravity conditions. It is conceivable that slower amyloid growth under microgravity conditions could also be attributed to suppression of the convective agitation.

Kinetic differences may have resulted in the observed morphological differences in Aβ(1-40) fibrils (**Figure 4–6**). Namely, under the gravity-agitated condition, fibrils were formed by kinetically stable inter-protofilament interactions, while the slow but steady growth without the effects of convective instabilities could have promoted optimal intra- and inter-molecular interaction modes during fibril growth. This gave rise to unique fibrous species, i.e. Type-2 fibrils, which are characterized by more extensive inter-protofilament interactions in comparison with Type 1 and Type-G, judging by the cross-sectional area of their amyloid core structures. Additionally, both Aβ(1-40) fibrils grown on the ISS (Type-1 and Type-2) showed a tendency to adopt a more twisted structure, with a higher pitch than ground-derived Type-G fibrils (**Figure 6, Table 1**). This may be also due to the lack of convection flows under microgravity conditions. In addition, the sedimentation effects under gravity conditions plausibly limit fibril morphology because the nuclei or growing fibrils come into contact with each other or with the bottom surface of the plate.

This study demonstrates that the ISS provides an ideal experimental environment for studying amyloid formation mechanisms by eliminating the conventionally uncontrollable gravity factor. The comparative analyses of amyloid fibril structures formed with and without gravity provide a deeper understanding of how microenvironmental factors surrounding assembling proteins affect fibril formation. Moreover, microgravity can offer unique opportunities for detailed observation of the earlier processes of amyloid formation, including nucleation and oligomer formation, since at least in Aβ(1-40), amyloid formation proceeds much more slowly in the absence of gravitational pull. By taking advantage of the microgravity experiments, systematic high-resolution analyses using isoforms and hereditary variants of Aβ are ongoing in our collaborative project. This series of studies will provide fundamental insights into the molecular mechanisms underlying amyloid formation and, more generally, the self-organization of biomacromolecules on Earth.

## Methods

### Sample preparation

Synthetic Aβ(1-40) was purchased from Toray Research Chemicals Co., Ltd.. Aβ(1-40) powder was dissolved at a concentration of 2 mM in 0.1% (v/v) ammonia solution, and diluted to a concentration of 0.1 mM with 20 mM sodium phosphate buffer (pH 8.0) containing 0.2 mM EDTA. Each sample was dispensed at 100 μL per well into multiple wells (0.2 ml each) of a 96-well PCR plate (semi-skirted with upstand, 4titue^R^). The top opening of the plate was sealed with a lid (Strips of 8 Flat Optical Caps, 4titude^R^) and covered with Kapton^R^ tape (DuPont-Toray Co., Ltd.). Sample preparation was performed on ice to prevent the formation of amyloid fibrils. The samples were frozen at −30°C for 3 h, and then frozen at −80°C for 16 h. Four plates were placed in one aluminum bag and sealed by Kapton^R^ tape. Totally 8 bags (8 sets of samples) were prepared and stored at −80°C until use.

### Procedures for space experiments on ISS-KIBO

For the spaceflight experiments, four of eight bags (four sets of samples) maintained at −80°C were launched by Space X-13 and transported to the ISS on December 16, 2017, and kept at - 95°C in the laboratory freezer (MELFI) in the Japanese experiment module KIBO (**Figure 2**).

All four sets of samples were stored in another MELFI set for 16 h at 2°C for defrosting, and then operated at 37±0.5°C to initiate the amyloid fibril formation in the Cell Biology Experiment Facility (CBEF) set. The sets of samples were taken out from the CBEF and immediately placed into MELFI at −95°C after incubating for 6 hours, 1, 3, and 9 days to stop further fibril formation. The samples were kept at −95°C in MELFI until just before the Space X-14 Dragon vehicle departed. The frozen samples landed on the Pacific Ocean on January 14, 2018 using the Dragon capsule, was transported to ExCELLS (Okazaki, Japan) and stored at −80°C until the experimental observations. A series of control experiments using the remaining four sets of samples were performed independently on the ground.

### ThT assay

For the ThT assay, three samples from each set, which had been incubated at 37°C for 6 hours, 1, 3, and 9-days and then frozen, were stored at 4°C for 17 h for thawing. Each sample solution was then dispensed at 100 μL per well with 200 μM of ThT into multiple wells of a clear-bottomed 96-well low-binding polyethylene glycol coating plate (Corning #3881). The sample plate was sealed with aluminum microplate sealing tape (Corning #6570). Fluorescence was measured immediately from the bottom of the plate at the excitation and emission wavelengths of 446 nm and 490 nm, respectively, by placing the 96-well plate at 25°C under quiescent conditions in a plate reader (TECAN).

### Cryo-EM

The Aβ(1-40) amyloid fibrils (0.1 mM) were diluted five times with 20 mM sodium phosphate buffer (pH 8.0) containing 0.2 mM EDTA and subjected to cryo-EM. A 2.5-μL aliquot was placed on an R 1.2/1.3 Quantifoil grid (Quantifoil Micro Tools) pre-treated by a glow discharge. Plunge-freezing of the specimen was performed at 4°C and 95% humidity using Vitrobot Mark-IV (Thermo Fisher Scientific). The frozen grids were kept in a cryo-storage under liquid nitrogen until use. For data collection, the grid was loaded into a JEM2200FS electron microscope equipped with a 200-kV field emission electron source using a Gatan 626 cryo-specimen holder at liquid nitrogen temperature. EM images were acquired on a DE20 direct electron detector (Direct Electron LP) at a nominal magnification of 30,000×, corresponding to 1.99 Å per pixel on the specimen. The images were corrected for beam-induced motion with dose-weighting using MotionCor2 software^32^ in the Relion 3.0 software^33^ and their contrast transfer functions were estimated using the CTFFIND4 software^34^. Micrographs were subjected to filament picking using e2helixboxer.py^35^ after reducing the image size by two binning. Filament images were extracted in 128 x 128 squares overlapping adjacent boxes by 90%. After 2D classification, the good classes, containing 1145, 404, and 774 segments of Type-1, Type-2, and Type-G fibrils, respectively, were used to generate each 3D model in Relion by assuming helical symmetry. The details are summarized in Table 1.

### Data Availability

The datasets generated and analyzed during the current study are available from the corresponding author on request.

## Acknowledgements

The Amyloid experiment was carried out through close collaborations among the Japan Aerospace Exploration Agency (JAXA), the Japan Space Forum (JSF), the National Aeronautics and Space Administration (NASA), and several related organizations and companies. We would like to thank our colleagues who were involved in this project for their efforts during this experiment. We are also grateful to Dr. Norishige Kanai (JAXA) and the ISS crew members for performing the experimental operations onboard the ISS. We thank Ms. Yukiko Isono (IMS) and Ms. Kumiko Hattori (Nagoya City University) for their help in sample preparation. We also thank the Biomolecular Dynamics Observation Group, ExCELLS, for the atomic force microscopy measurements.

This work was supported in part by JSPS KAKENHI Grant Numbers JP17K15441 and JP19K07041 to M.Y-U. This work was also supported by the Cooperative Study Program (19-261) of National Institute for Physiological Sciences.

## Author contributions

M. Y-U., S. Y., C. Y., H. K., T. Shimazu, K. K. designed the experiments.

M. Y-U., S. Y., C. Y., H. K., T. Shimazu carried out the sample preparation.

M. Y-U. and S. Y. carried out the kinetics experiment on the ground.

M. Y-U., C. S. and K. M. carried out the cryo-EM experiments.

M. Y-U., S. Y. and T. Satoh analyzed the kinetic and cryo-EM data.

M. Y-U. and K. K. wrote the paper.

## Competing interests

The authors declare no conflict of interest.

